# Hyperspectral Remote Sensing for Harmful Algal Bloom Detection: *Pseudo-nitzschia* in the Northeast Pacific

**DOI:** 10.64898/2026.02.24.707776

**Authors:** Alexander L. Bailess, Nicholas Baetge, Andrew H. Barnard, Nicholas Tufillaro, Michael Behrenfeld, Brian Bill, Raphael Kudela, Jason Graff, Maria Kavanaugh

## Abstract

Diatoms are microscopic marine algae that are critical for global primary production, carbon sequestration, and fisheries productivity. However, select diatoms may form harmful algal blooms, which threaten marine ecosystems and the fisheries they sustain. Rapidly identifying harmful blooms is necessary to effectively manage marine resources, yet current identification methods are limited by expensive and labor-intensive *in situ* point sampling.

Hyperspectral remote sensing enables scalable monitoring, but its ability to resolve taxonomic shifts within phytoplankton groups (e.g. diatoms) is largely unknown. To investigate this uncertainty, we cultured four dominant diatom genera from the California Current upwelling system, including this systems’ most abundant harmful algae, *Pseudo-nitzschia*. The hyperspectral absorption and backscatter of these taxa were measured and used to model spectral reflectances that remote sensing platforms (satellites/drones) might detect. Differences between fingerprints of these taxa were quantified using vector-based and statistical analyses.

Mean spectral differences of 48% were observed between the most dominant diatom, *Thalassiosira*, and the most toxic diatom, *Pseudo-nitzschia*. Differences of approximately 30% were found between *Pseudo-nitzschia* and the second and third most abundant diatoms, *Chaetoceros* and *Asterionellopsis*. Successful identification of *Pseudo-nitzschia’s* reflectance fingerprint was driven by the presence of a unique feature around 560 nm. The distinct spectral fingerprint of *Pseudo-nitzschia* indicates that it can be distinguished from benign diatom blooms using hyperspectral remote sensing platforms.

## 2 Introduction

Diatoms are an important subgroup of phytoplankton, comprising 10 and 20% of global and marine primary production, respectively (Behrenfeld et al. 2021, Tréguer et al. 1995, Moore et al. 2001b, Aumont et al. 2003), and up to 40% of the marine biological pump (Jin et al. 2006, Tréguer et al. 2018). Large-bodied diatoms also support healthy fisheries by mediating efficient energy transfer to higher trophic levels (Chavez, Messié, and Pennington 2011). Of the 1000 plus known genera of diatoms (Fourtanier and Kociolek 2003), most facilitate beneficial ecosystem services. However, a few diatoms can be harmful, producing toxins or physical structures detrimental to fisheries and human health.

Harmful algal blooms (HABs) are aggregations of algae that threaten human health, ecosystems, or human activities (Kudela et al. 2015). In up-welling systems, some of the most toxic HABs come from the genus *Pseudo-nitzschia*. Half of *Pseudo-nitzschia* species produce a potent neurotoxin, domoic acid, that accumulates in primary consumers (e.g. bivalves, zooplankton, small fish) before being concentrated in marine mammals, seabirds, and fishes of commercial importance. For marine mammals and humans, domoic acid poisoning is devastating, causing the swelling and death of neurons, which induces disorientation, seizures, temporary or permanent memory loss, and in some cases, coma and death (Bates et al. 2018).

The world’s largest fisheries are found within the highly productive Eastern Boundary Upwelling Systems (Ryther 1969, Chavez and Messié 2009). Here, large-scale nutrient injection supports plankton blooms, which ultimately sustain these fisheries. However, these blooms can be episodically dominated by toxigenic *Pseudo-nitzschia*, with devastating ecological and economic effects—for example the coastwide *Pseudo-nitzschia* bloom in 2015 that spread from Baja California to British Columbia (McCabe et al. 2016). This bloom closed recreational and commercial fisheries, including mussels, razor clams, oysters, Dungeness and rock crabs, anchovies, and sardines. The closures resulted in an estimated revenue loss of $170 million for the Dungeness Crab fishery alone (Pacific States Marine Fisheries Dungeness Crab Report 2014) (inflation adjusted to ∼$230 million in 2025), placing stress on fishing communities and coastal towns (Moore et al. 2020). The high levels of domoic acid caused severe damage to marine ecosystems with effects propagating throughout the food web. Mass mortalities and strandings of tens of thousands of seabirds, seals, and sea lions occurred, with toxins also detected in whales and dolphins (McCabe et al. 2016, Gibble et al. 2018).

Accurate and timely detection of large *Pseudo-nitzschia* blooms is critical for fisheries management, aquaculture operations, and public health. Current monitoring methods leverage discrete samples of domoic acid accumulation in shellfish tissues with microscopy-based identification of *Pseudo-nitzschia* cells. While these methods provide essential information, they suffer from limited spatial and temporal resolution and delayed detection relative to contamination (Lefebvre et al. 2017). Annual costs of this continuous *in-situ* monitoring are estimated in the millions of dollars (Anderson et al. 2000). These challenges underscore the need for contemporary methods—such as emerging remote sensing approaches—that can accurately detect *Pseudo-nitzschia* blooms across broader spatial and temporal scales.

Harmful algal bloom identification is the first step in ecological analysis, fisheries management, and early warning response. Identification using ocean color is likely the most cost-effective way to assess ecosystem threats over large spatial and temporal scales. To identify HABs using ocean color, we must first develop spectral ‘fingerprints’. Fingerprints are derived by measuring the optical properties (absorption and backscatter) of a specific taxon which, when combined (equation 1), mirror the signal an ocean color remote sensing platform will observe (Gordon et al. 1988, Zaneveld 1995). In practice, isolating fingerprints from the ocean is often confounded by the overlapping optical properties of dissolved substances, detritus, and other phytoplankton. However, the bloom-forming diatoms cultured in this study form dense surface aggregations *in-situ* (Trainer et al. 2009) that dominate the ocean color signal. This dominance makes them particularly well-suited for identification using spectral fingerprints. Moreover, with the launch of NASA’s latest satellite mission, PACE (Werdell et al. 2019), we now have unprecedented spectral coverage of the global oceans. This hyperspectral resolution provides significantly more information about the phytoplankton within the water, illuminating features and subtleties previously unseen.

Satellite-measured ocean color is strongly influenced by particulate backscatter. Backscattered light is all light that returns in the direction of its source after interaction with a particle or surface. The physical properties of this particle or surface (shape, size, and refractive index) influences the intensity, geometry, and spectral characteristics of the backscattered light (Boss et al. 2004). Unlike absorption spectra, whose features are easily associated to their physical drivers (e.g. pigments), the relationships between backscatter spectral features and their physical drivers are more nuanced. Consequently, there is limited understanding of how hyperspectral backscatter’s features relate to their physical drivers. Historically, partly due to limited spectral resolution, spectral backscatter was assumed to be a monotonic function, without pronounced spectral features (e.g. Lee, Carder, and Arnone 2002, Lee et al. 2009, Kostadinov, Siegel, and Maritorena 2009, Morel and Prieur 1977). While this assumption works well for clear-oceanic waters (Smith and Baker 1981), coastal and upwelling systems with non log-linear particle size distributions (e.g. plankton blooms) and non-spherical particle shapes violate the basis for this assumption. Indeed, hyperspectral backscatter contains substantially more diagnostic information than spectral slope alone, with unique and more pronounced spectral features than the spectrum of total scatter (this study, Bricaud, Morel, and Prieur 1983, and Morel and Bricaud 1981). The advent of new hyperspectral backscatter technologies (e.g. Novak, Burmeister, and Röttgers 2024) will provide a wealth of new data to link *in situ* biogeochemistry and community composition (including HAB dominance) to hyperspectral backscatter features—which may be derived from NASA PACE. This will advance ocean color inversion models significantly, increasing their utility in detecting HABs by incorporating the information from empirically derived backscatter spectral shapes.

## 3 Methods

### 3.1 Cultures

Absorption and backscatter spectra were measured from acclimated cultures of four diatom genera, known to be the most dominant within the California Current. They are, in order of typical *in-situ* biovolume: *Thalassiosira rotula, Chaetoceros affinis, Asterionellopsis glacialis*, and *Pseudo-nitzschia spp*. (Du, Peterson, and O’Higgins 2015, Lassiter et al. 2006). Isolates were obtained from the Bigelow National Center for Marine Algae, except for *Pseudo-nitzschia*, which were clonal isolates from the Washington and Oregon coast (*P. fraudulenta, P. pungens*, and *P. seriata*). Each taxon was kept in exponential growth in triplicate 2 L Nalgene bottles as semicontinuous batch cultures. All cultures were grown in an environmental control room at 15°C with a salinity of 33 PSU, using L1 media (Guillard and Hargraves 1993). Cultures were exposed to 400 *µmol* photons *m*^2^/s with a relatively flat, white spectrum. The intensity of the grow lights (a Phyton-Systems light bank) were measured using a QSL-scalar PAR sensor. This light bank was scheduled on a 12:12 sinusoidal light-dark cycle to simulate a natural day-night cycle. Cultures were kept in suspension by bubbling with two aquarium air pumps (ProFILE 5500). Pumped air was filtered through a hydrocarbon trap (Restek) before entering the cultures.

Cells were acclimated to growth conditions (light, nutrients, temperature) for two months (*>>* 30 divisions) before experimental sampling to ensure steady-state, photoacclimated growth. Cell and detrital abundance were quantified daily via an Imaging Flow CytoBot (IFCB) to derive growth rates and ensure detrital production remained low. Using light microscopy, the presence of detritus too small for the IFCB was observed, and quantified using a Coulter Counter (Beckman Multisizer 3, 70 *µ*m aperture). The biomass of detritus was small compared to that of the live cells and detrital particles were almost entirely *<* 3 *µ*m, much smaller than this study’s phytoplankton which were *>* 10 *µ*m (Table 1). Phytoplankton photophysiology was monitored using a fast repetition rate fluorometer (FRRf). Fast repetition rate fluorometry samples were exposed to 15 *µmol* photons/*m*^2^/s at 480 nm for 10 minutes to relax non-photochemical quenching (NPQ) before measurements were made (Milligan, Aparicio, and Behrenfeld 2012). The amount of time and total irradiance to fully relax NPQ varied by taxon and time of day. We conducted several experiments and found 15 *µmol* photons/*m*^2^/s for 10 minutes to yield the highest Fv/Fm (∼0.6 ± 0.05) for these taxa on average; Fv/Fm values *>* 0.55 indicated healthy diatoms ready for inline system measurements (Tan et al. 2019, Dijkman and Kromkamp 2006).

**Table 1.** Approximate cell size ranges for representative diatom species.

| Species | Individual Cell Sizes |  |
| --- | --- | --- |
| | width<br>($\mu\text{m}$) | length<br>($\mu\text{m}$) |
| *Thalassiosira rotula* | 8 - 60 | 5 - 20 |
| *Chaetoceros affinis* | 8 - 15 | 7 - 30 |
| *Asterionellopsis glacialis* | 5 - 8 | 30 - 150 |
| *Pseudo-nitzschia* spp. | 3 - 8 | 70 - 145 |

### 3.2 Inline system

Measurements of phytoplankton absorption, scatter, and attenuation coefficients were made using a SeaBird (formerly WETLabs) AC-S (4 nm spectral sampling frequency). Backscatter was measured using a Sequoia Hyper-BB (5 nm spectral sampling frequency). All bio-optical measurements were made following the best practices for IOP measurements (Boss et al. 2019). Instruments were configured in a darkened recirculating flow-through inline system driven by a peristaltic pump (Slade et al. 2010). The inline system was filled with 33 PSU, 0.2 *µm* filtered seawater, leaving sufficient headspace to add 200mL of phytoplankton culture. After inoculation, the solution was allowed to recirculate and homogenize until AC-S and HyperBB data approached a stable asymptote. Measurements were recorded for 10 to 15 minutes and the median derived for each wavelength to reduce the influence of outliers and noise. This process was repeated using a 5.0 *µ*m filtrate of the sample to estimate the optical contributions of detritus and colored dissolved organic matter (CDOM) to the culture. Subtraction of the 5 *µ*m filtrate’s optical signature from that of the total culture isolated the diatom-specific optical signature.

Between daily sampling, the inline system was flushed with deionized water. The optical windows and tubes of the AC-S were cleaned with 98% isopropyl alcohol and Milli-Q water, while the plastic HyperBB windows were cleaned with soap and rinsed with Milli-Q. After cleaning, degassed Milli-Q blanks were recorded for both the AC-S and HyperBB, which were then taken apart and cleaned again until two baseline measurements agreed at the 0.005 m^-1^ level. All bio-optical data were recorded using the software Inlinino (https://github.com/OceanOptics/Inlinino, Haëntjens and Boss 2020).

Measurements of absorption, backscatter, and total scatter were recorded for triplicates of all genera, except *Asterionellopsis*. Due to a gain issue with the HyperBB, only a single backscatter spectrum was obtained for *Thalassiosira* and due to a pump failure at the end of sampling, the backscattering coefficients for the final *Pseudo-nitzschia* triplicate, PN 3, were measured by placing this sample directly into the HyperBB light trap. The solution was stirred thoroughly to homogenize and then measured (this sample had no detrital blank). To align the spectral sampling frequency (4 nm for the AC-S and 5 nm for the HyperBB) and center wavelengths of both instruments, ACS data were linearly interpolated to 1 nm resolution and then down-sampled to 5 nm. Intrataxon backscatter and absorption data were then bootstrapped to provide more spectra for clustering algorithm analysis. All backscatter spectra from triplicate cultures were divided by all absorption spectra of the same genera (e.g. the backscatter spectra of PN 1 was divided by absorption of PN 1, 2, and 3. The same was done for the backscatter spectra of PN 2 and so on). The primary clustering algorithm used was an unsupervised machine learning algorithm, k-means, from the open-source Python machine learning library Scikit-learn. The optimal number of clusters (k) was determined using an elbow method and silhouette scores. To isolate differences in spectral shape, independent of magnitude, a mean-normalization was performed where each value within a spectrum was divided by the mean of the entire spectrum.

### 3.3 Modeling

The spectral fingerprints of our *in vivo* cultures, measured using the inline system, can be related to *in situ* reflectances using the following equation (Gordon et al. 1988, Zaneveld 1995):

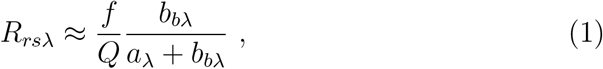

where *R*_*rs*_(reflectance) describes the intensity and color of light, at wavelength *λ*, leaving the ocean’s surface. The backscatter coefficient at wavelength *λ* is represented by *b*_*b(λ)*_, while *a*_*(λ)*_ denotes the absorption coefficient at wavelength *λ*. The symbols *f* and *Q* represent scaling factors which do not affect the spectral shape. Because *a*_*(λ)*_ is generally multiple orders of magnitude greater than *b*_*b(λ)*_, we can further approximate the equation to:

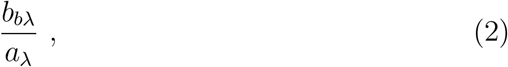

as the contribution of backscatter in the denominator is often negligible.

### 3.4 Distance Metrics

Distance functions were used to distinguish between modeled spectral fingerprints in a reliable and quantitative way. These functions are able to define the degree of similarity between complex, non-linear spectra by assessing differences in translation (center wavelength or hue), and standard deviation (full-width half max (FWHM) or peakedness), of spectral features. Deborah, Richard, and Hardeberg 2015 and Deborah 2016 evaluated thirty-one distance functions for hyperspectral analysis. Of these, we chose the Spectral Correlation Mapper (SCM) (Carvalho Júnior and Meneses 2000) as the optimal tool for calculating the differences between phytoplankton fingerprints. Spectral Correlation Mapper builds from vector-based angular methods which have a history of successful applications in remote sensing (e.g. Wei et al. 2022, Kruse et al. 1993). These tools are unaffected by changes in spectral magnitude and provide absolute differences between spectra. Spectral Correlation Mapper uses Pearson’s correlation coefficient (r):

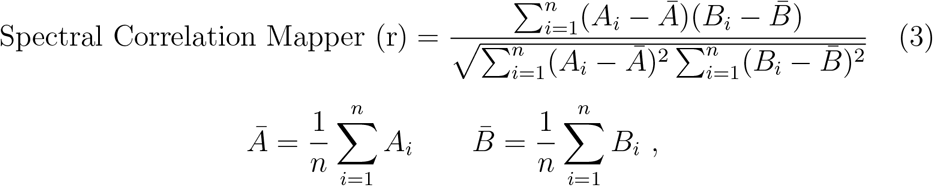

to evaluate the degree of positive or negative linear correlation between mean-standardized data. Spectral Correlation Mapper processes spectra as a distribution, appropriate for the highly correlated nature of hyperspectral data.

Spectral Correlation Mapper’s sensitivity to differences between spectral signatures can be further improved through the application of derivative transformations. However, while derivatives excel at highlighting small features, the increased sensitivity also amplifies noise, which must first be removed from the spectra. To remove noise while preserving as many of the spectral features as possible, we used a Savitzky-Golay filter. The Savitzky-Golay method preserves large spectral features better than other techniques (such as mean or median filtering) by fitting an ‘x’ order polynomial to the data within the filter window. This preserves spectrally-wide nonlinear features while spectrally-narrow variations (noise) are smoothed out. For these reasons, Savitzky-Golay smoothing is often used in hyperspectral applications (e.g. Vandermeulen et al. 2017, Xi et al. 2015, Hunt 2024). The Savitzky-Golay filter has two tunable parameters, window length and polynomial order. The goal is to find the ‘goldilocks’ parameters where the data are neither oversmoothed (thus losing important information) nor overfit (turning noise into features). We used the Scipy.signal function savgol filter and chose a window length of 13 (5 nm spacing x 13 = 65 nm windows) and a polynomial order of 4 based on the iterations seen in the supplemental materials (figure 11). Savgol filter ‘mode’ was set to ‘nearest’ to reduce boundary artifacts from the filter window interacting with the end of the spectra (mode ‘nearest’ pads the end with values identical to the last encountered). The option ‘deriv’ was set to 2 (takes the second derivative of the spectra).

## 4 Results

All cultured phytoplankton share local maxima of backscatter around 560/570 nm (figure 2), the yellow-green region of these spectra. These backscatter maxima are likely amplified by the intersection of absorbance features, from fucoxanthin and the secondary peaks of chlorophylls b and c, which form a local absorbance minima (figure 3). However, the absolute lowest absorbance for all species throughout most of the visible range lies near 600 nm. While we observe local maxima in backscatter at 600 nm (figure 2), these wavelengths do not correspond to the greatest backscatter for any species. Instead, local and total maxima around 560/570 nm appear in nearly all species, being most pronounced for *Asterionellopsis* and *Pseudo-nitzschia*. For *Pseudo-nitzschia*, the 560/570 nm peak is followed by a steep decline that exhibits a subtle inflection near 585 nm before decreasing further. This subtle inflection coupled with the aforementioned peak in *Pseudo-nitzschia’s* backscatter spectra creates an important feature in *Pseudo-nitzschia’s* reflectance fingerprint, seen as a bifurcated peak in figure 4. After the maximum green-yellow (560/570 nm) backscatter peak, *Pseudo-nitzschia* and *Asterionellopsis* backscatter less for bluer wavelengths, contrary to *Thalassiosira* and *Chaetoceros* (figure 2). There is also a consistent trough for all spectra around 660 nm followed by a large peak that overlaps with the chlorophyll Qy absorption band around 670 nm (figure 3).

**Figure 1.**
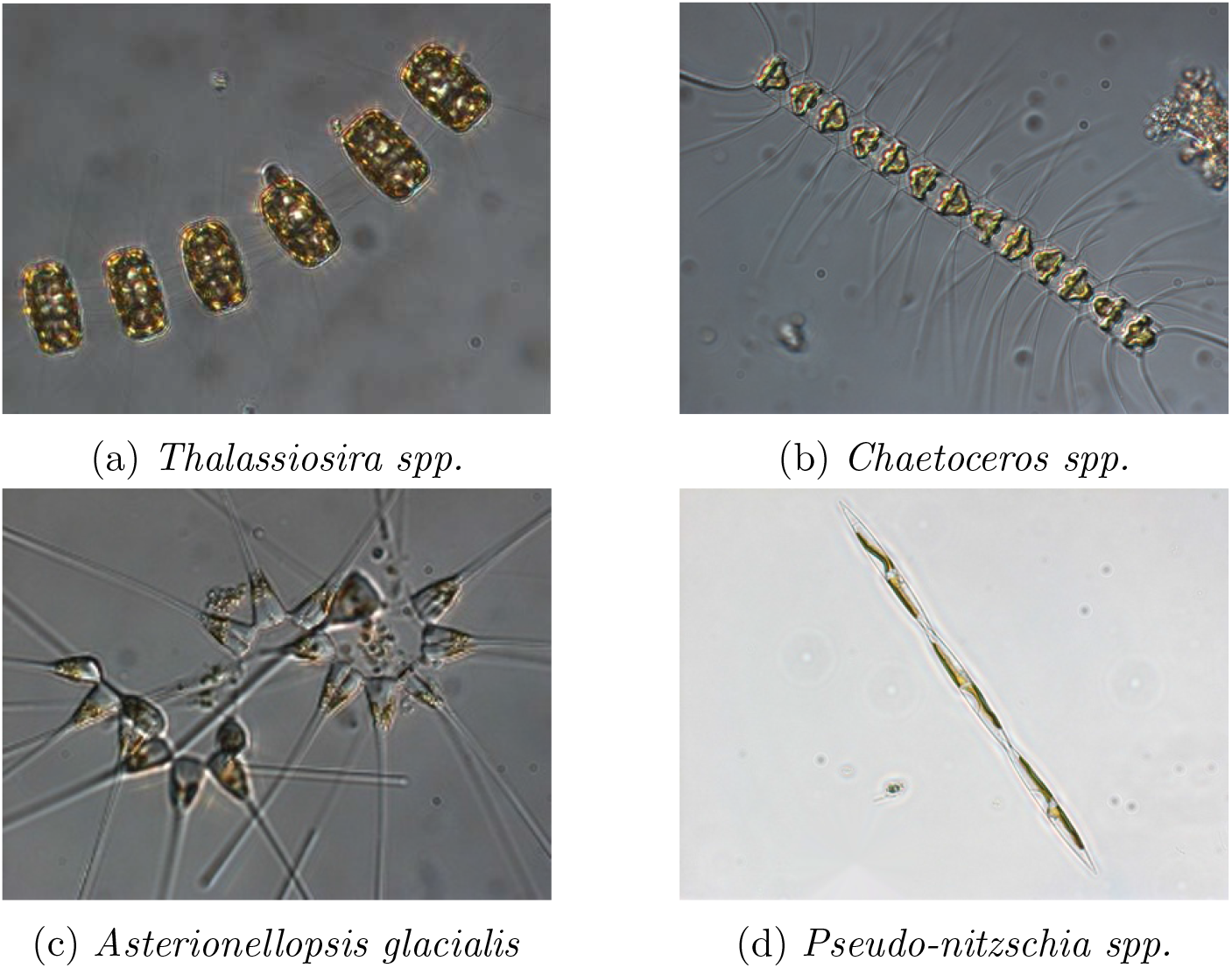
Representative diatom genera (reproduced with permission from the Kudela Lab).

**Figure 2.**
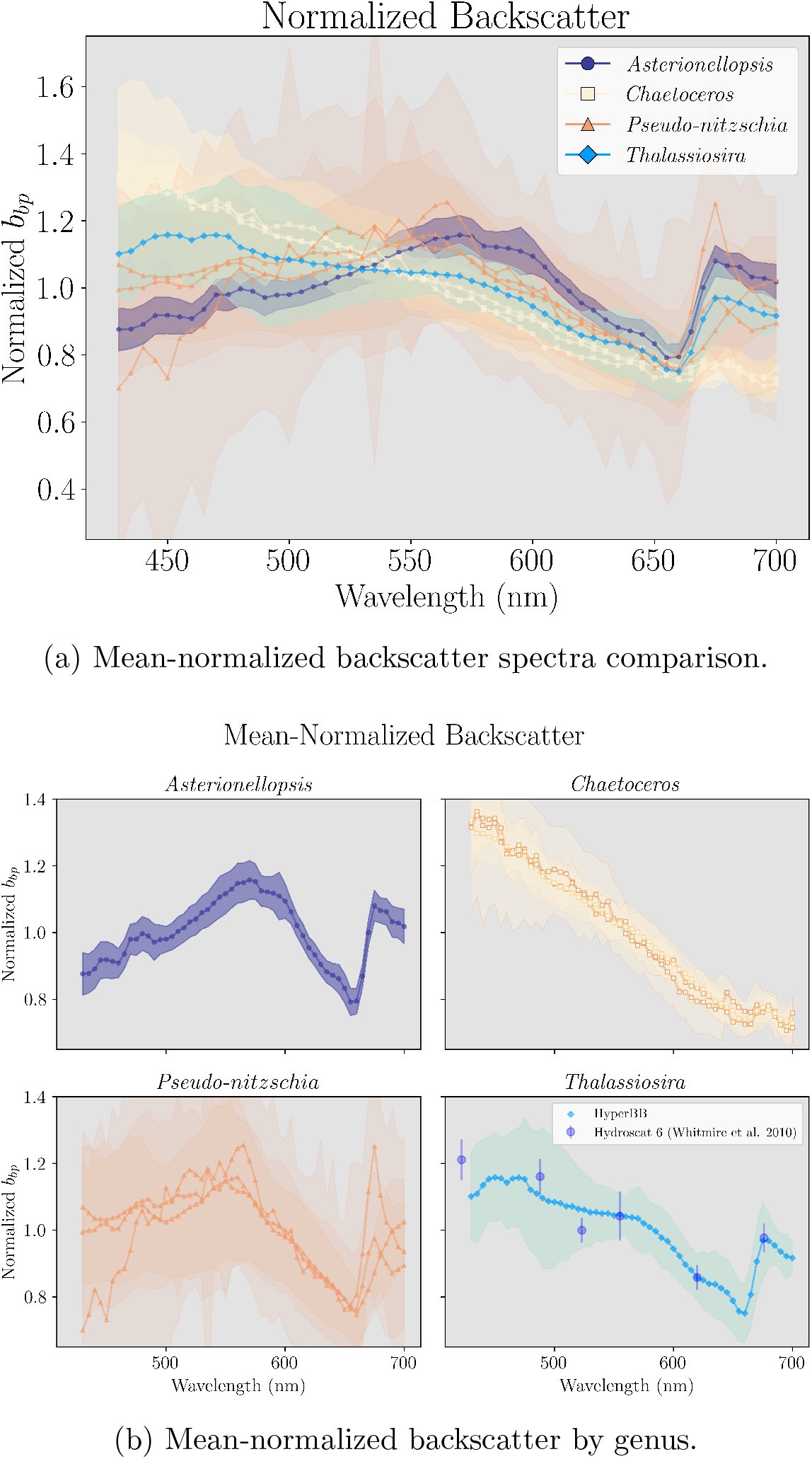
Normalized backscatter spectra for each species. Dots represent the normalized median value of backscatter at each wavelength measured. Ribbons represent one standard deviation from the median, colors distinguish each species. Triplicate samples were measured for *Pseudo-nitzschia* and *Chaetoceros*. A multispectral comparison of *Thalassiosira rotula* from Whitmire et al. 2010 is included in dark blue, with bars representing 1 standard deviation from the median. Multispectral and hyperspectral data were aligned at 620 nm for comparison.

**Figure 3.**
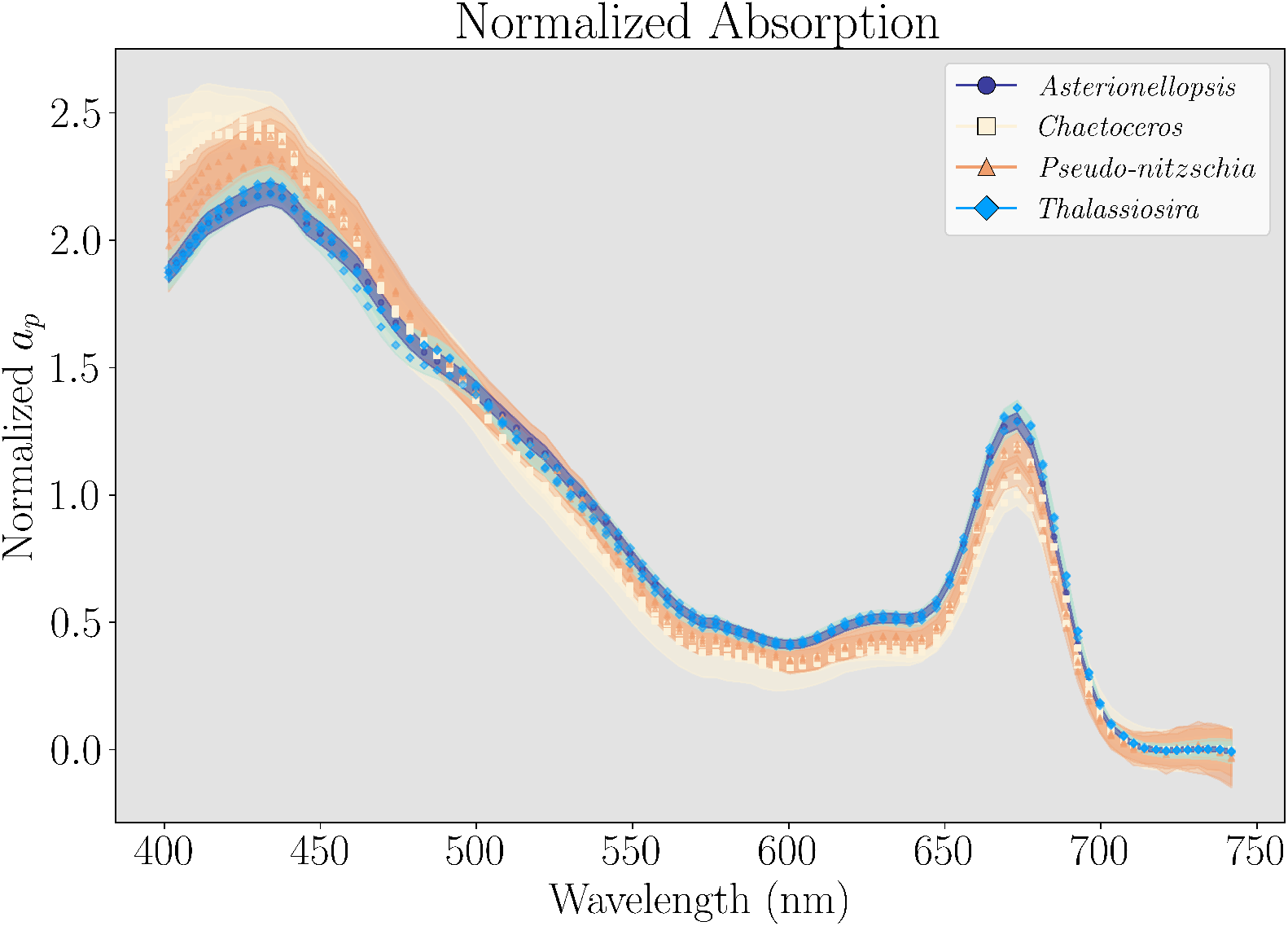
All cultures absorption spectra using the AC-s, spectra are mean-normalized. The peak centered on 440 nm is the Soret band, and the peak over 670 nm is the Qy band.

**Figure 4.**
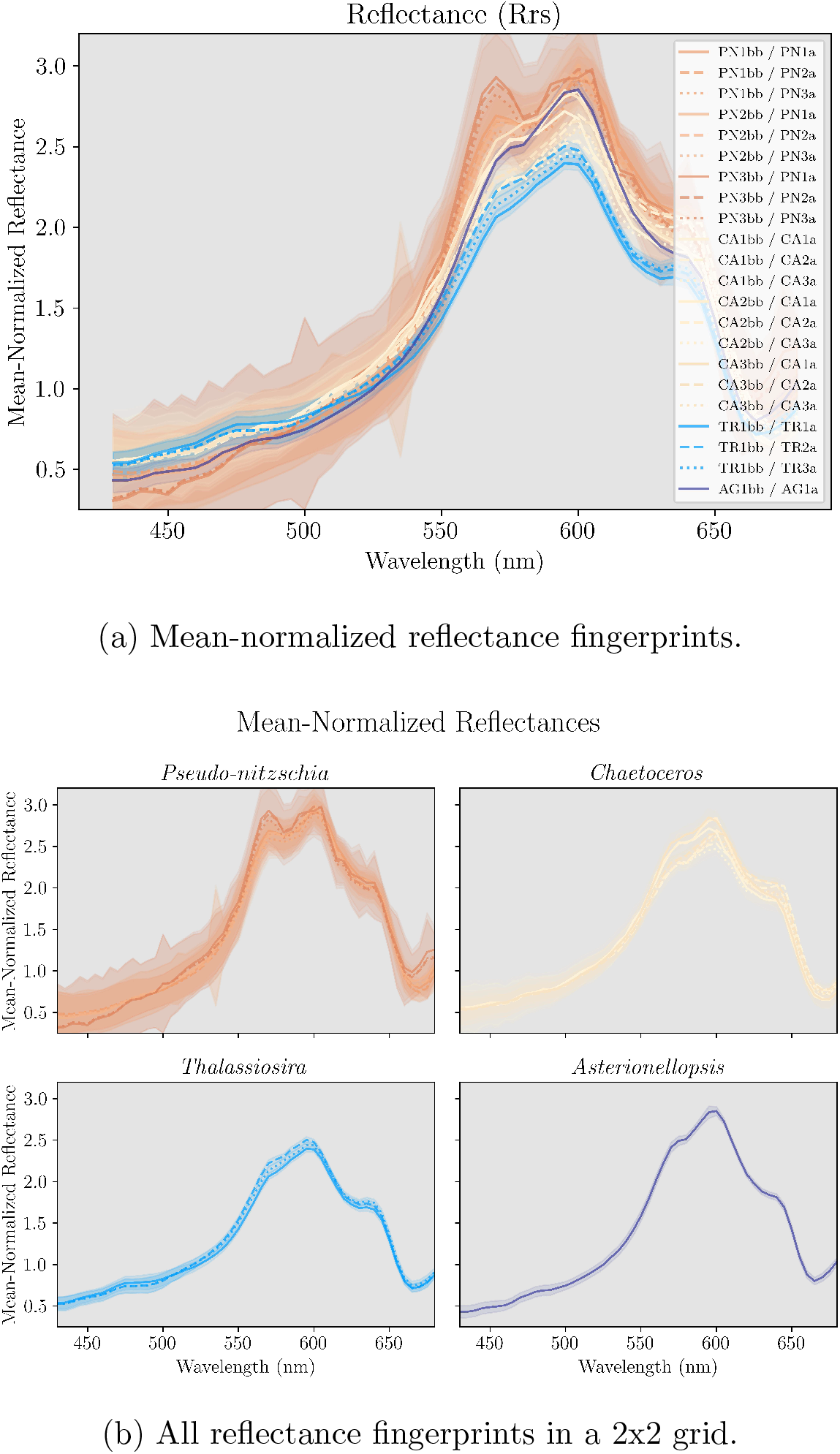
Nineteen bootstrapped spectra of the four diatom genera measured in this experiment: *Asterionellopsis* (AG) is purple, *Chaetoceros* (CA) is yellow, *Pseudo-nitzschia* (PN) is orange, and *Thalassiosira* (TR) is blue. Each spectrum is the mean value of hundreds to thousands of absorption and backscatter measurements (for each *λ*_*i*_), converted to reflectance spectra using equation 2.

Figure 3 shows the absorption spectra for each of the cultured diatoms. Triplicate cultures within each taxon exhibit consistent spectral shapes across wavelengths. Differences in the absorption spectra between taxa are less visually pronounced than those observed in normalized backscatter (figure 2) or total scatter spectra (figure 9, supplemental material). There are small deviations in the absorption spectra of the third triplicate sample of *Pseudonitzschia* (PN 3) from the first and second replicates (PN 1, PN 2), with slightly elevated mean-normalized values in the blue region (400 – 440 nm), likely related to the aforementioned pump failure for this sample. *Thalassiosira* has the strongest shoulder at 470 nm from the pigment fucoxanthin. *Asterionellopsis* has the largest ratio of Qy to Soret band absorption (chlorophyll a absorption in the red vs blue, respectively). *Chaetoceros* appears to have the broadest peak for the Soret band and the lowest relative absorption of carotenoids (pigments in roughly the 470 - 650 nm range).

The most pronounced distinction in the spectral shapes of reflectance (figure 4) is for *Pseudo-nitzschia*, which forms distinct bifurcated peaks around 570 and 600 nm, with a trough at 585 nm. This feature provides robust separation of the spectral fingerprint of *Pseudo-nitzschia* from all other diatoms sampled. In contrast, *Chaetoceros* forms a shoulder near 570 before peaking around 600 nm, while *Thalassiosira* remains linear throughout this range, monotonically increasing. The spectral fingerprint of *Asterionellopsis* is similar to that of *Chaetoceros*, with a shoulder at 570 nm and a peak at 600 nm, however the height of the peak relative to the shoulder is greater for *Asterionellopsis*. For all genera, there are only small differences in slope from the deep-blue to cyan (∼430 to 510 nm), with the largest differences in yellow and orange (∼560 to 620 nm). There is relative uniformity at all other wavelengths. Because diatom absorption spectra are highly similar, genus-level discrimination is primarily driven by spectral backscatter. Unique backscatter features are most visible in reflectance spectra when centered within the absorbance minima of 550 to 650 nm.

Unsupervised machine learning algorithms were able to successfully distinguish between the spectral fingerprints of *Pseudo-nitzschia, Chaetoceros*, and *Thalassiosira* (figure 5). The highest silhouette coefficient found used k=5 (0.5), but k=2-4 were all within 8% of this score (0.46 - 0.48). Using k=2 (clustering the spectra into two groups) isolates the *Pseudo-nitzschia* spectra, separating them from all other taxa. Using k=3 provides nearly perfect clustering of spectra, with the exception that *Asterionellopsis* is misclassified as *Chaetoceros*. Using k=4 yields similar results, however *Pseudo-nitzschia* is divided into two clusters. Lastly, using k=5 divides *Chaetoceros* into two clusters (CA1bb triplicates and AG, versus CA2bb and CA3bb triplicates). Other non-Euclidean machine learning algorithms were applied and results from these are provided in the appendix.

**Figure 5.**
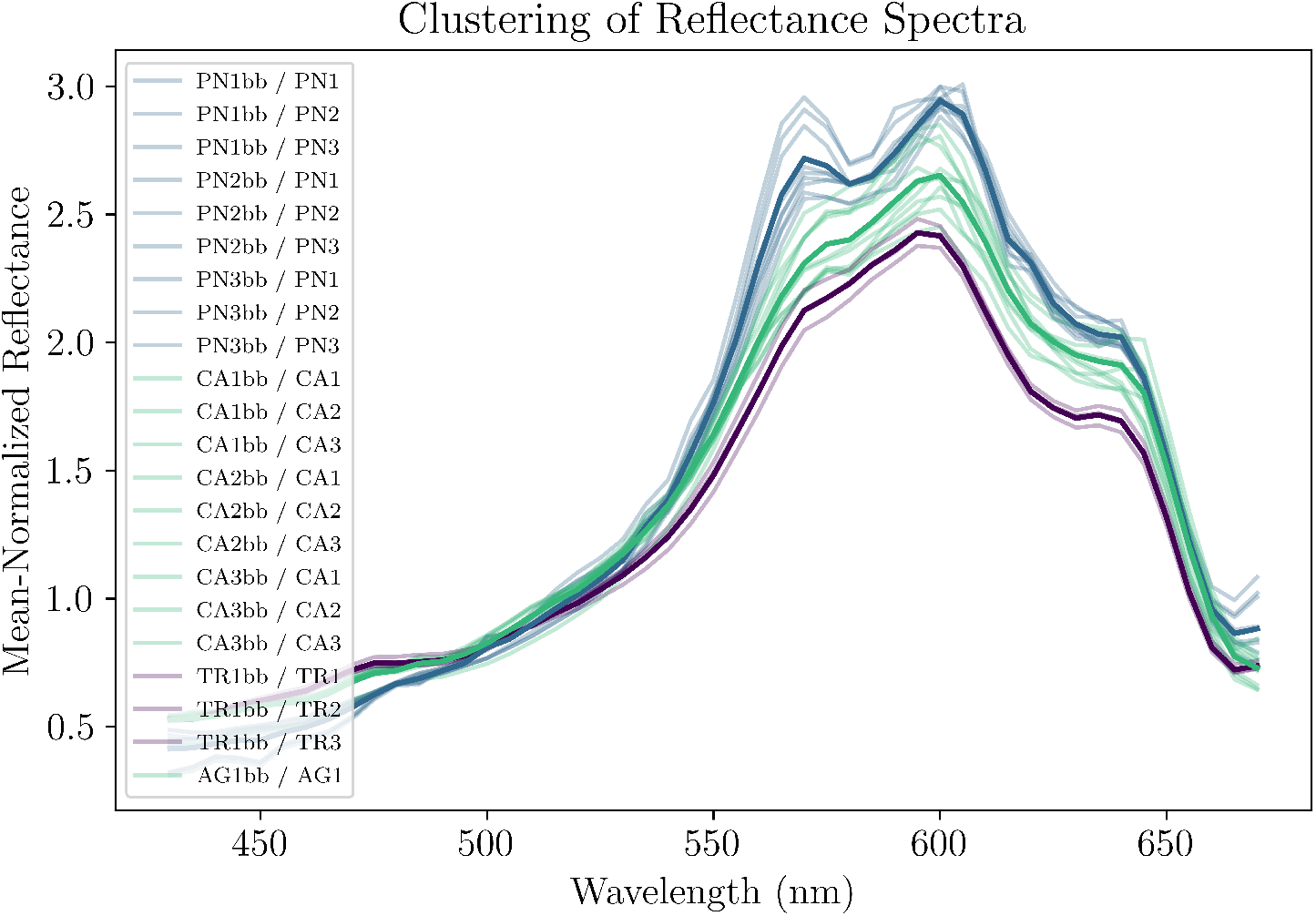
All bootstrapped spectra clustered using the k-means algorithm for k=3. *Pseudo-nitzschia* is abbreviated as ‘PN’, *Chaetoceros* as ‘CA’, Thalassiosira as ‘TR’, and Asterionellopsis as ‘AG’. The number following the two letter phytoplankton ID code corresponds to the replicate culture number. ‘bb’ refers to backscatter. Therefore, **PN1bb/PN2** corresponds to the backscatter spectra of the first replicate culture of *Pseudo-nitzschia* divided by the absorption spectra of the second replicate culture of *Pseudo-nitzschia* (a bootstrapped sample). Measured spectra are shown semi-transparent while the derived mean of each cluster is opaque. Ignoring *Asterionellopsis* (only one sample), the clustering algorithm successfully classified all taxa.

To accentuate spectral features, the second derivatives of reflectance fingerprints from figure 4 were calculated and are shown in figure 6. These smoothed, second derivative fingerprints were then compared using the Spectral Correlation Mapper (SCM) distance metric. A heatmap of the evaluated differences is shown in figure 7. The SCM performed better when subsetting the spectra in figure 6 from 550 to 600 nm, wavelengths where the majority of spectral differences are found. Spectral Correlation Mapper calculates all *Pseudo-nitzschia* spectra as having a mean intertaxon correlation of 98% with a standard deviation of ± 1% (upper-left, red, right triangle), while *Thalassiosira* spectra are deemed nearly identical to each other (99% or greater correlation). *Chaetoceros* spectra are the most variable with a standard deviation of 7% and an average correlation of 91%. Spectral correlation mapper finds average differences of 29% ± 13% between *Pseudo-nitzschia* and *Chaetoceros* (top, black box), and 34% ± 7% between *Pseudo-nitzschia* and *Asterionellopsis* (bottom, black rectangle). Spectral correlation mapper performed exceedingly well for *Pseudo-nitzschia* and *Thalassiosira* (middle, black rectangle)—the most toxic and most common diatoms, respectively— with average differences of 48% ± 10%.

**Figure 6.**
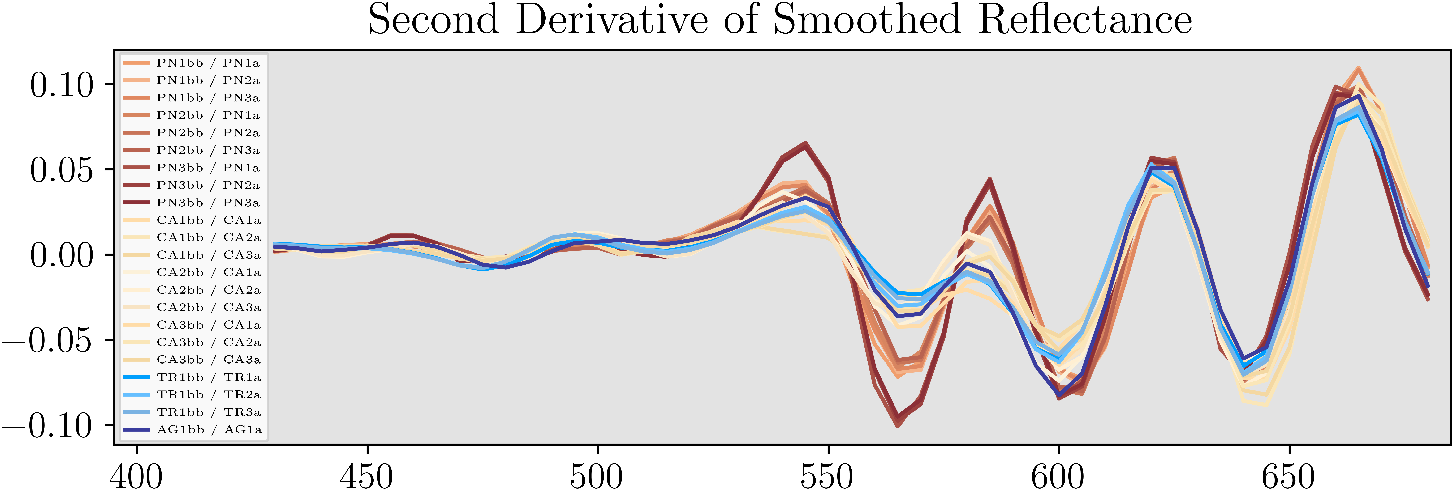
Second derivative of the smoothed spectra.

**Figure 7.**
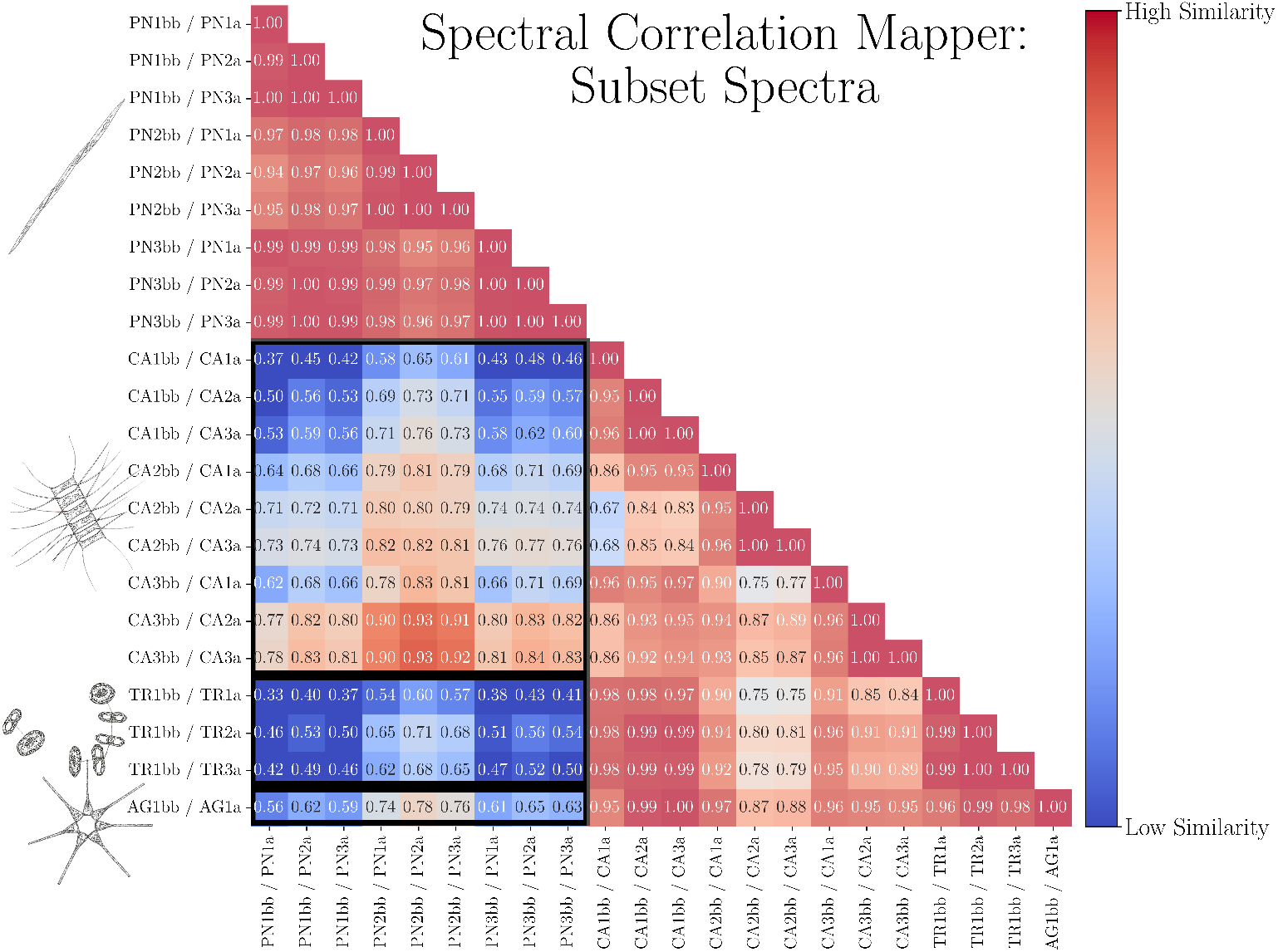
Spectral Correlation Mapper applied to a subset of the reflectance spectra (550 to 600 nm) from figure 6 to focus on regions of maximum differences. The black boxes highlight comparisons between the fingerprints of *Pseudo-nitzschia* (PN) and *Chaetoceros* (CA), *Thalassiosira* (TR), and *Asterionellopsis* (AG), in descending order—summarized in table 2.

## 5 Discussion

Our approach yielded distinct reflectance fingerprints, each related to a unique phytoplankton genera under bloom conditions. We identified visually discernible, group-specific spectral features and quantified these differences. *Pseudo-nitzschia*, the dominant HAB taxa, exhibits a unique feature in its modeled reflectance fingerprint, a bifurcated peak between 570 and 600 nm. This feature easily distinguishes the spectral fingerprint of *Pseudo-nitzschia* from those of the three most abundant diatoms in this system—both through visual assessment and using unsupervised classifying algorithms. *Pseudonitzschia*’s fingerprint was most easily distinguished from *Thalassiosira’s*, which was nearly flat between 565 and 600 nm. This distinction is important because *Thalassiosisra* is the most abundant bloom-forming diatom in the Northern California Current. The ability to discern between *Thalassiosira* and *Pseudo-nitzschia* based on spectral fingerprints is valuable for a number of applications, such as productivity modeling, species distribution mapping, and HAB monitoring. While *Chaetoceros* fingerprints were more variable between these wavelengths, visual assessments and clustering algorithms were still able to distinguish between the fingerprint of *Chaetoceros*, the second most abundant diatom, and *Pseudo-nitzschia*.

Unsupervised clustering algorithms (figure 5 and appendix, figure 10) successfully isolated the unique spectral fingerprints of *Pseudo-nitzschia* from those of *Thalassiosira*, and *Chaetoceros*. These algorithms proved robust and versatile, successfully classifying *Pseudo-nitzschia’s* fingerprint regardless of spectral permutations (full spectral range or subset spectra, raw spectra or the second derivative, smoothed spectra or unsmoothed). Clustering algorithm’s success with these fingerprints suggests an automated spectral classification framework will be able to distinguish between monospecific or near monospecific blooms of the most common diatoms and *Pseudo-nitzschia* using the PACE satellite.

### 5.1 Importance of backscatter

These experiments reveal the importance of integrating measured spectral backscatter into ocean color models. Rethinking the assumption that spectral backscatter is relatively featureless, being either flat or a monotonic function, may help us to disentangle the complexity of case 2 waters (optically-complex waters influenced by terrigenous inputs, namely CDOM and sediments). Under the assumption of spectrally flat backscatter, reflectances are merely the inverse of absorption (figure 8). Spectral shapes in these ‘pseudoreflectances’ are nearly identical, with no translational differences between spectral features and only minor changes in the FWHM of peaks at 600 nm. Meanwhile, the variability and features seen in figure 4 highlight the complexity of parameters that define the spectra of backscatter: unique shapes, sizes, and intracellular structures. It is due to the unique morphological features, stemming from the great diversity of the diatoms, that the complexity in the spectral shapes of backscatter—and consequently, reflectance fingerprints— enables the identification of *Pseudo-nitzschia* from other diatom assemblages.

**Figure 8.**
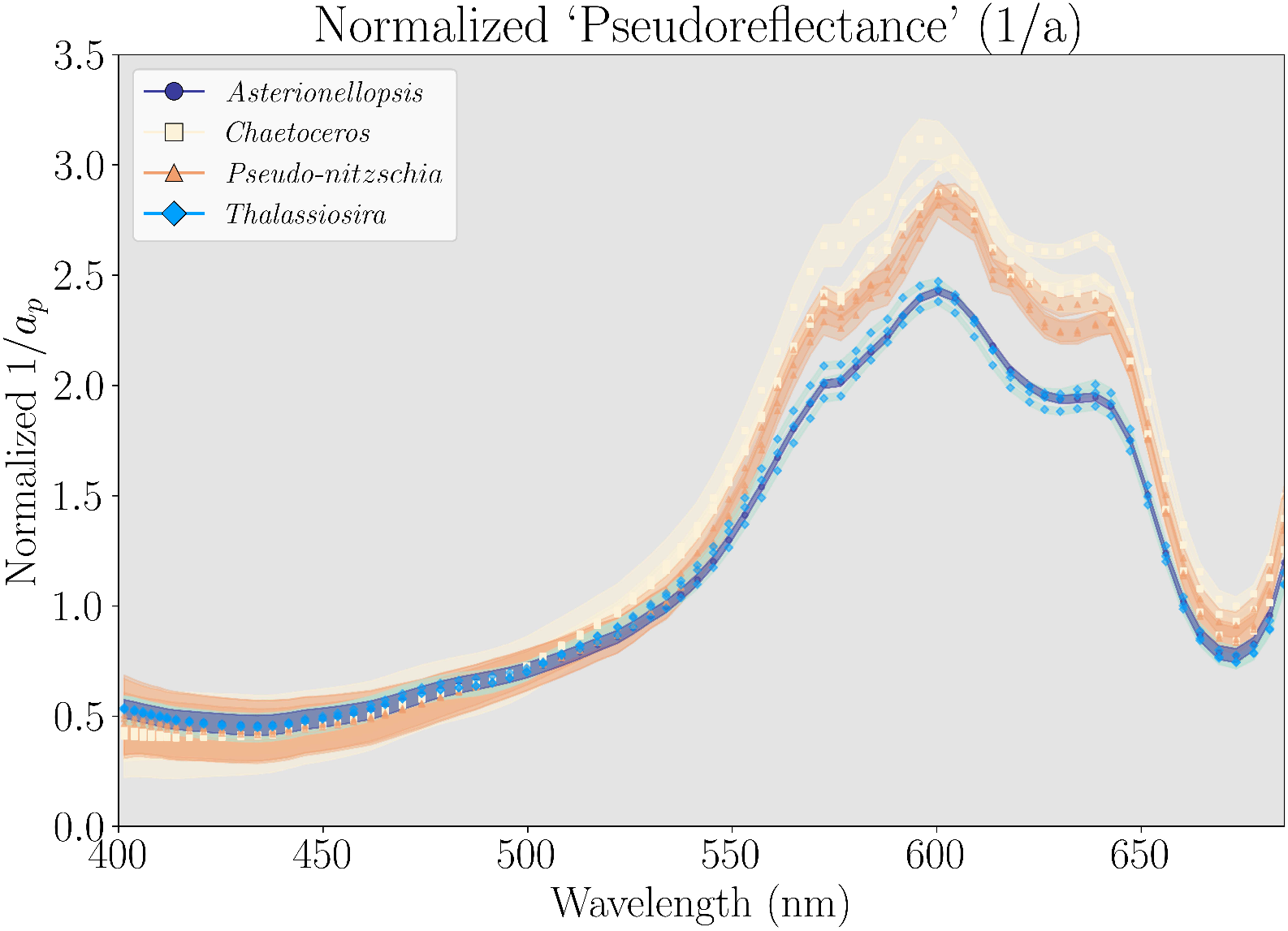
What if backscatter was flat? Pseudoreflectances shown assuming backscatter is spectrally flat for all samples.

Our results indicate that absorption-based models will be unable to discern *Pseudo-nitzschia* in mixed diatom assemblages, while reflectance-based models that integrate spectral backscatter have potential. This is because spectral backscatter presents unique features and shapes (figure 2). Modeling the backscatter of these diatom’s complex frustule geometries and intracellular structures is prohibitively computationally intensive, making direct attribution of individual spectral features to morphologies challenging. However, we hypothesize that the continuous increase of backscatter coefficients in bluer wavelengths for *Chaetoceros* is due to the presence of many thin setae (spines) whose small size should preferentially scatter shorter (bluer) wavelengths of light. A similar increase in backscattering coefficients of bluer wavelengths is observed for *Thalassiosira*, due to the presence of many small chitinous thread structures which adorn the frustule (Tran et al. 2023). Many of these threads are joined together in the center of the frustule to bind *Thalassiosira* cells in chain formations. We propose that chitins’ increased absorption in blue-green wavelengths (Azofeifa, Arguedas, and Vargas 2012), relative to silica, makes these thread structures less efficient backscatters than setae. The varying efficiencies of backscatter for these distal structures may explain the observed differences in figure 2a between *Thalassiosira* and *Chaetoceros* in blue-green wavelengths. We note that natural populations of *Asterionellopsis glacialis* often have large siliceous spikes, similar to *Chaetoceros* setae, however in our cultures the length of these spikes were reduced compared to natural samples. This may explain why *Asterionellopsis* does not have the same high backscatter coefficients in blue wavelengths as *Chaetoceros* and *Thalassiosira*. However, both *Chaetoceros* and *Thalassiosira* possess numerous setae or threads per cell, whereas *Asterionellopsis* has but a single, large spike per individual. As a result, the backscatter spectrum of *Asterionellopsis* in blue-green wavelengths may be more strongly influenced by absorption features—the Soret band and fucoxanthin—than by elevated backscatter from a single siliceous spike, leading to the spectral shape observed in figure 2.

The peak in backscatter for all species past 660 nm (figure 2a) is likely due to inelastic scatter in the form of fluorescence, rather than true backscatter. This fluorescence signal results from the large FWHM (∽25 nm) of light emitted and accepted by the HyperBB in red wavelengths. Past 660 nm, the Hyperbb efficiently excites phytoplankton photosystems due to a high absorption coefficient from the chlorophyll Qy band. Some of the absorbed photons are subsequently re-emitted by the phytoplankton at slightly redder wavelengths as fluorescence, which are then detected by the instrument and registered as backscatter. This effect appears to be an instrument-specific artifact of the HyperBB’s detection window, rather than a property of phytoplankton scattering itself. Nevertheless, agreeing with the postulates of Bricaud, Morel, and Prieur, 1983, the backscatter spectra presented more spectral complexity and shape, and therefore information, than the spectra of total scatter alone (figure 2 versus supplemental material, figure 9).

### 5.2 Orientation of cells

During these experiments, diatoms were measured in a turbulent environment that randomized their orientation. Since these taxa have complex, non-spherical shapes, stochastic orientation creates a plethora of apparent sizes and shapes (cross-sectional areas and geometries, relative to the incoming light source and detector). These apparent shapes and sizes will affect the optical properties of the cells (Marcos et al. 2011).

We observed an increase in the variability of backscatter coefficients with bluer wavelengths in all taxa, but especially *Pseudo-nitzschia*. Mie theory— which can be used to derive the optical properties of a single, homogeneous sphere—offers an explanation, citing backscatter coefficients of bluer wavelengths to be more sensitive to changes in apparent size (Mie 1908, Bohren and Huffman 2004, Van de Hulst 2012). Mie’s derivations are based on spherical particles, which do not adequately represent the complex morphologies of diatoms examined here. However, backscatter coefficients and spectral features for complex (non-spherical) particles are poorly constrained (Clavano, Boss, and Karp-Boss 2007), making Mie theory an imperfect but indispensable framework for inference in particle-light interactions.

Hence, we theorize that the large variability in backscatter coefficients for bluer wavelengths is due to changes in apparent size. This is supported by *Pseudo-nitzschia*’s backscatter spectra, which possesses the greatest variability of all backscatter spectra, especially for bluer wavelengths. Accordingly, *Pseudo-nitzschia* cells present the largest differences in apparent size between the apical axis and top-down, valve-apex view. The apical axis presents a large cross-sectional area when facing the light-source and detector, an oblate spheroid. While in a top-down view of the valve-apex, even *Pseudo-nitzschia* forming long chains present but a small, nearly spheroid, cross-sectional-area to the sensor. This small, spherical cross section would elevate backscatter in bluer wavelengths, while reducing overall scatter (Mie 1908, Marcos et al. 2011). The difference in apparent size between apical and valve-apex views of *Pseudo-nitzschia* is greater than that of any other diatom sampled in this experiment and will increase multiplicatively with chain length. Indeed, due to *Pseudo-nitzschia*’s unique chain morphology, this difference in apparent size, and the resulting variability in backscatter coefficients, is likely higher than the vast majority of other marine phytoplankton, excluding a few penates with similar morphologies and chain-formation patterns (e.g. *Rhizosolenia*). In a turbulent medium (e.g. our inline system) where orientation is randomized, the direct valve-apex view will be significantly less probable than some angle of the planar view.

In their natural environment, cells are often assumed to be well mixed in the euphotic zone, but this is not always the case. During periods of laminar flow or stratification, some phytoplankton will orient themselves to the currents or sunlight, respectively (Karp-Boss and Jumars 1998, Nayak et al. 2018). Aligned orientation alters the apparent cross-sectional area of cells and can substantially modify both the magnitude (sometimes by more than 30% (Marcos et al. 2011)) and spectral characteristics of backscattered light. The optical effects of orientation will be most strongly expressed in the polarization of backscatter. Multi-angle polarimetry from platforms such as PACE offers a way to assess orientation-dependent scattering and polarization in natural assemblages, which will help to constrain uncertainties associated with phytoplankton taxonomic retrievals.

### 5.3 Distance metrics and proposed use

To quantify differences between spectra, we used the distance function SCM. Compared to other distance functions, SCM is easy to interpret (Pearsons’ correlation coefficient (r)) and insensitive to changes in magnitude, while still responsive to differences in spectral shape (Carvalho Júnior and Meneses 2000). Spectral Correlation Mapper performed best after applying the second derivative to the reflectance fingerprints as spectral features were accentuated. This quantification of spectral differences is important for establishing baselines to compare ocean color data to modeled reflectance fingerprints.

The low standard deviation between mean reflectance fingerprints of *Pseudonitzschia* replicates (1%, figure 7) suggests that stringent cutoffs can be applied when looking for *Pseudo-nitzschia*-dominated blooms. Differences between the most abundant diatoms and *Pseudo-nitzschia* ranged from 29 to 48% (table 2). Thus, when comparing ocean color data to the modeled fingerprints of *Pseudo-nitzschia*, the threshold for *similarity* between *in-situ* and modeled spectra should be greater than 71%. Ocean color datasets can be flagged for pixels with high spectral similarity to the modeled reflectance fingerprints, providing an automated and scalable framework.

**Table 2.** Mean spectral differences between *Pseudo-nitzschia* and other genera, with standard deviations.

| Mean Difference and Standard Deviation of Spectral Fingerprints:<br>Comparing *Pseudo-Nitzschia* and other Diatoms |  |
| --- | --- |
| Taxa | *Pseudo-nitzschia* spp. |
| *Chaetoceros affinis* | 29% ± 13% |
| *Thalassiosira rotula* | 48% ± 10% |
| *Asterionellopsis glacialis* | 34% ± 7% |

These distance metrics perform best under bloom-conditions when the optical properties of the dominant phytoplankton are the primary contributor to ocean color. Removing the reflectance spectrum of pure water from the ocean color signal should produce a spectrum that closely matches the modeled fingerprints—if the bloom is dominated by one of these modeled taxa.

However, there will be greater spectral variation in natural samples due to mixed assemblages and other optically active components (colored dissolved organic matter (CDOM), detritus, sediments). This variability will make it difficult for distance metrics to select pixels where key phytoplankton groups are present, as the influence of other optically active components will shift the spectral shape further from that of the modeled fingerprints. Fortunately, the spectral shapes of CDOM and detritus are generally featureless, monotonic functions in visible wavelengths (Aurin, Mannino, and Lary 2018). Additionally, while the optical signatures of CDOM and detritus can have significant influence on the magnitude of ultraviolet and blue wavelengths, these spectral regions are ignored in this proposed framework for diatom identification. Corrections can also be made to account for CDOM contributions using widely-accepted remote sensing algorithms (e.g. Zhu et al. 2011, Aurin, Mannino, and Lary 2018, Bélanger, Babin, and Larouche 2008, Lee et al. 2009). The resuspension of sediments in the nearshore environment will be more difficult to account for, as the reflectance of these materials align with the yellow-orange wavelengths (560 - 630 nm) used for *Pseudo-nitzschia* identification. This may prevent identification in turbid estuaries and the very immediate nearshore environment. However, these spaces account for a small fraction of the total area within upwelling systems.

The primary obstacle in diatom identification will be the competing optical properties of mixed assemblages. While distance metrics should still be used to pull out pixels (spatial bins) with high similarity to modeled fingerprints, this should be followed up with further analysis into the spectra of pixels that do not pass the similarity threshold. This can be done by first using a clustering algorithm to subset the region of interest into ‘k’ bins with similar spectral shapes. From these k-bins, a visual analysis or band ratios near 570 and 585 nm, may be used to identify *Pseudo-nitzschia* within mixed assemblages. The competing optical properties of many unique phytoplankton taxa may denature the structure of the *in-situ* reflectance too much for distance metrics to discern. However, if *Pseudo-nitzschia* is present in sufficient concentration, the distinct ‘M’-shape seen in figure 4 should be present.

Inverse models and hyperspectral libraries will benefit from future studies investigating the detection limits of unique taxa within mixed community assemblages, as well as additional measurements of taxa-specific hyperspectral backscatter. As higher spectral resolution and lower bandpass sensors become available, specifically open-source and accessible tools (e.g. Novak, Burmeister, and Röttgers 2024), we anticipate other genus-specific spectral features to become known. This will further constrain uncertainties in ocean color models including the model proposed in this study.

### 5.4 Integration with Existing Monitoring Frameworks

The California—Harmful Algae Risk Mapping (C-HARM) model was developed to combat the uncertainty in *Pseudo-nitzschia*’s presence and variable toxin production along the California coast (Anderson et al. 2016, 2011, 2009, 2006). California—Harmful Algae Risk Mapping integrates regional ocean circulation and statistical models with remote sensing data to provide forecasted predictions for the presence of toxic *Pseudo-nitzschia*. The remote sensing data integrated into the model predicts *Pseudo-nitzschia* cellular abundance using the ratio of reflectance from 488 to 555 nm, while the intensity of reflectance at 555 nm is coupled with reflectance-derived chlorophyll and other products to estimate toxin concentration. These remote sensing products and bands, while useful correlative variables within the Southern California Current, are not strictly tied to *Pseudo-nitzschia*. Hence, extrapolation to higher latitudes within the California Current may be less reliable if the biogeochemical relationships to *Pseudo-nitzschia* and its toxicity are dynamic.

The PACE satellite’s hyperspectral resolution will allow us to better target the spectral regions of interest identified above—providing an optical proxy *directly tied* to the inherent optical properties of *Pseudo-nitzschia* itself. We anticipate that these findings can augment the current C-HARM model by integrating hyperspectral genus-specific fingerprinting using PACE data. This may increase C-HARM’s skill when extrapolating to new environments, e.g. the Northern California Current (Oregon, Washington, British Columbia). This is especially important because these more temperate zones must be sampled for *Pseudo-nitzschia* and domoic acid with greater frequency than Central and Southern California, due to strong seasonality and rapid bloom progression (Frolov, Kudela, and Bellingham 2013). This direct spectral pathway for detection may further allow the extrapolation of this approach to other systems where *Pseudo-nitzschia* is present in harmful concentrations.

### 5.5 Conclusion

In this study, we measured the hyperspectral backscatter coefficients of the dominant diatom genera within the California Current, including the region’s most prominent HAB genera, *Pseudo-nitzschia*. Hyperspectral reflectances were then modeled and spectral differences derived through quantitative comparison. Our results support future algorithm development using remote sensing for the identification of harmful *Pseudo-nitzschia* blooms, particularly using hyperspectral observations from missions such as NASA PACE. This capability may be used to inform local agencies of HAB events in near-real time and facilitate targeted *in situ* sampling of toxin levels.

While additional field-based studies will be necessary to validate the fingerprints derived from these experiments, our data provide evidence that significant differences exist in the spectral shapes of these key plankton that are sufficient to discern between them using the methods provided. These results offer a new approach for monitoring biodiversity and detecting harmful algal blooms.

## 5.6 Acknowledgments

We would like to thank all students and staff of the 2023 Ocean Optics Class and the 2022 Inline Workshop at the University of Maine. These courses fostered a close-knit community, provided mentorship on best practices in ocean optics, and introduced students to valuable open-source resources such as *Inlinino*. Emmanuel Boss, Collin Roesler, Meg Estapa, Ivona Cetinić, Wayne Slade, Nils Haëntjens, Patrick Gray, Charlotte Begouen Demeaux, and Guillaume Bourdin lead lectures and labs, giving students the knowledge that made this work possible. We are also indebted to many for conversations and advice that greatly benefited this work, including Mallie Hunt, Tara Conrad, Kim Halsey, Allen Milligan, Robert Frouin, Ashley Ohall, and Luke Carberry. We are deeply grateful to friends who helped us with sampling, setup, and late-night culture maintenance: Kelly George, Finley Dibert-Burgwin, Wave Moretto, and Ian Black. We are grateful for support from the OSU Seascape Ecology Lab, OSU Phytoplankton Ecophysiology Lab, and resources from the UCSC Kudela Lab, and the OSU Giovannoni lab.

## 6 Supplemental Materials

**Additional Distance Metrics** Additional distance metrics explored included: cosine similarity, Euclidean distance of cumulative spectra, and Kullback-Leibler pseudo-divergence.

Cosine similarity is defined as:

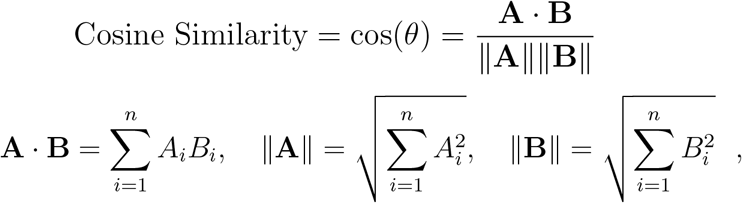

where *A*_*i*_ and *B*_*i*_ are two distinct spectra. Cosine similarity takes one point of a spectrum (i) and places it in one-dimensional space (a floating dot), it then iterates upon this, placing another point (i+1) and adding a dimension (d+1). We now have two coordinates in 2-dimensional space to which we can draw a line from the origin (i.e. a vector in an x - y plane). The algorithm continues to add coordinates and subsequent dimensions until we have a space with equal dimensions and coordinates as our spectrum (in this case we are evaluating wavelengths 430 - 670 at 5 nm increments (i & d = 48)). For these spectra we create a single vector in 48-dimensional space derived from the coordinates and take the cosine of this vector, spectrum a, to that derived for spectrum b; where spectra a and b represent the ‘fingerprints’ of two unique phytoplankton taxa. When we apply a weighting function to the spectra (to isolate colors with the most inter-taxa variability), we reduce the dimensionality to the number of wavelengths being analyzed. It should be noted that cosine similarity is processed in Euclidean space and therefore assumes different wavelengths are statistically independent from one another. We know of course that this is untrue, as the distinct bands that compose hyperspectral data are highly correlated (440 nm is deep blue, and 441 nm is just slightly less deep blue).

In cosine space when two vectors are orthogonal to each other (at a 90^°^angle), we infer that they are entirely different by the metrics that cosine similarity measures. In contrast, vectors with a 0^°^ angle indicate identical spectra. Cosine similarity is a non-linear metric, for example, a cosine value of 0.9 corresponds to an arccos(0.9) = 24.8^°^ angular difference in n-dimensional space (a 0.9 - 1 = 10% difference in cosine similarity space), while a value of 0.99 would be an arccos(0.99) = 8.1^°^ angular difference (0.99 - 1 = 1% difference in cosine similarity space).

Deborah 2016 found cosine similarity to work well with theoretical data but when applied to real spectra, noise contaminated accurate measurements. This issue can be alleviated by a Savitzky-Golay smoothing function. Both cosine similarity and SCM are common in the literature and were designed to ignore magnitude changes—we often only care about spectral shape and not intensity.

While cosine similarity and SCM respond well to overlapping translational changes (wavelength/hue changes), they saturates when these changes become too large (when spectral features no longer overlap). This is not an issue for our comparisons as the center wavelengths of spectral features are well-aligned, largely due to the common pigment classes shared between diatoms. Thus, the dominant parameter in our distance metric is the standard deviation of a spectral features, which cosine similarity and SCM excel at measuring.

The two following methods are sensitive to changes in magnitude and do not saturate with increased distance between features. These are the Euclidean distance of Cumulative Sum (ECS) and the Spectral KullbackLeibler Pseudo-Divergence (KLPD). Both of these metrics fully satisfy the three fundamental criterion (magnitude, translation, and standard deviation) for a distance metric. One caveat to ECS is that this function integrates the spectra from the lowest to highest values, placing more emphasis on differences in bluer wavelengths. The integration can be flipped to place more emphasis on the red wavelengths, however, the derived differences between spectra will change depending on how the metric is integrated. Kullback-Leibler pseudo-divergence is a variation on the standard Kullback-Leibler divergence function, which is used for probability density functions, hence *pseudo*-divergence. Kullback-Leibler pseudo-divergence violates the theory of triangular inequality which means that we expect different, albeit similar, responses when comparing spectrum i to spectrum j as when comparing spectrum j to spectrum i. However, all spectral features are evenly weighted. For both ECS and KLPD, a value closer to 0 means the spectra are more similar, unlike cosine and SCM where a value closer to 0 means the spectra are more dissimilar.

Euclidean distance of Cumulative Spectra (ECS), processes the spectra as a distribution:

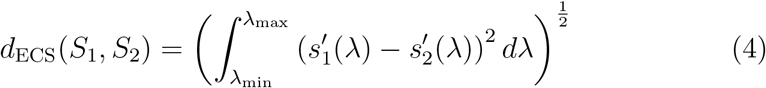

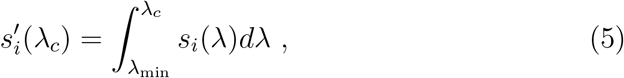

where *s*_1_ and *s*_2_ are two distinct spectra, or ‘fingerprints’ to be compared. Kullback-Leibler Pseudo-Divergence (KLPD) is similar to ECS, but is not spectrally weighted:

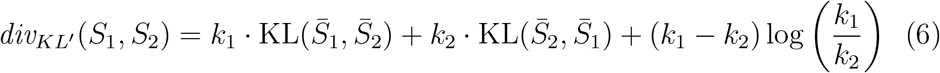

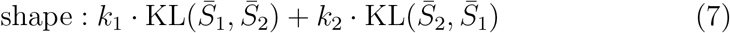

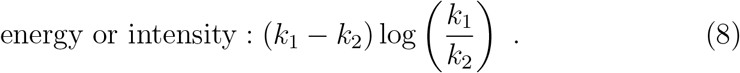

Both ECS and KLPD performed well (intertaxon differences were large, while intrataxon differences were small) but are only able to provide relative differences, rather than absolute values like cosine similarity and SCM.

The value in both ECS and KLPD comes from their ability to show the relative differences between fingerprints. Euclidean distance of Cumulative Spectra presents the largest dynamic range with the center, identical values (whited out) at 0 while maximum differences are interpreted for *Thalassiosira* and *Pseudo-nitzschia* around 850. KLPD presents lower dynamic range, reaching a maximum of 59 for *Thalassiosira* and *Pseudo-nitzschia*, but presents the same general patterns. Looking at figure 13 we can see three red hotspots; one in the top left, the center-bottom-right, and nearly the bottom right. These are emphasizing the intrataxon similarities between *Pseudo-nitzschia, Chaetoceros*, and *Thalassiosira*.

On the full raw spectra, cosine similarity performed poorly in terms of percentage differences, all spectra were given identical values to the 0.00 level (spectral differences were all *<* 1% in cosine space (heatmap not shown)). However, when we increased the decimal limit, we found that although differences were *<* 1%, these small differences were still sufficient to successfully cluster most of the reflectance fingerprints into their respective taxonomic groups (figure 10). Both ECS and KLPD were performed on the full raw spectra because they are able to measure differences in magnitude and do not saturate with distance for translational changes. Cosine similarity and SCM perform best after a derivative is applied and the spectra is subset to a region of interest.

**Figure 9.**
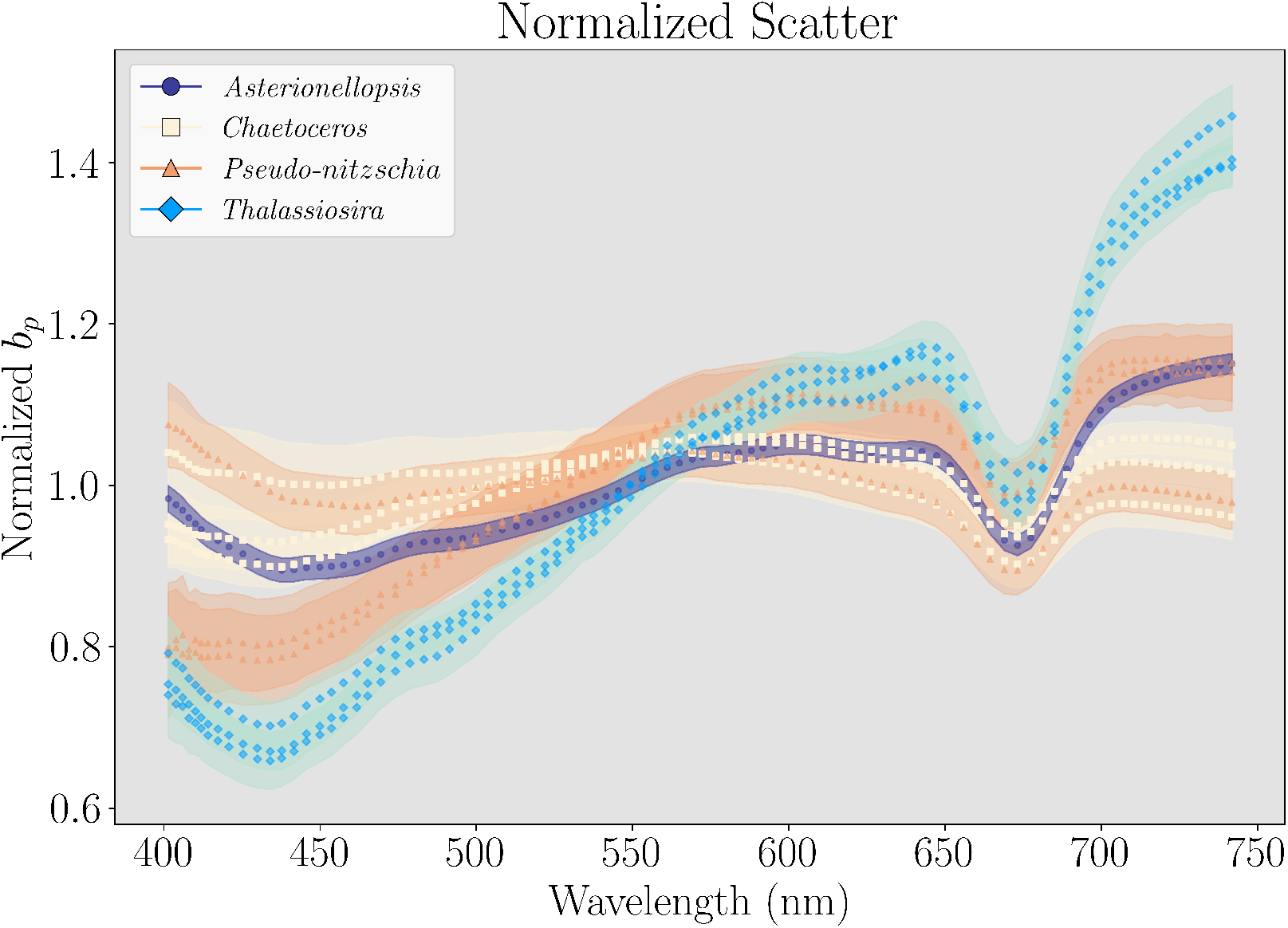
Normalized total scatter spectra for all species. All species have total scatter spectra that visually appear to be more similar intraspecifically than interspecifically. The standard deviations are low across all species. PN 3 is different from other triplicates due to a pump failure (the more flat, orange line). These spectra of total scatter closely mimic an inversion of the absorption spectra.

**Figure 10.**
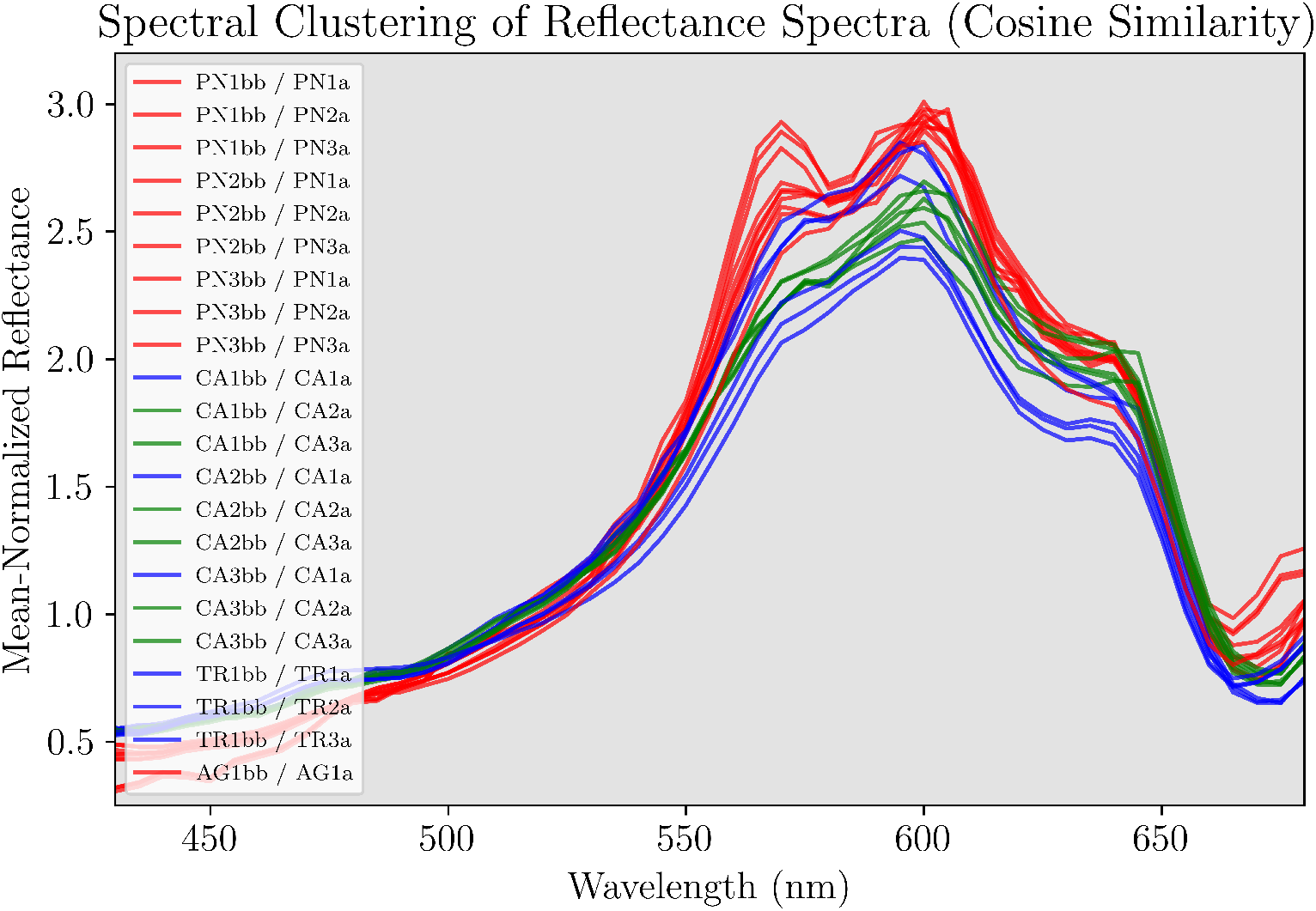
Spectral clustering: unsupervised machine learning using cosine similarity on all raw spectra. Cluster number set to 3. The absorption spectra of the first triplicate of Chaetoceros (CA1a) is misclassified as Thalassiosira (TR) for all bootstrapped spectra. Asterionellopsis is misclassified as Pseudonitzschia.

**Figure 11.**
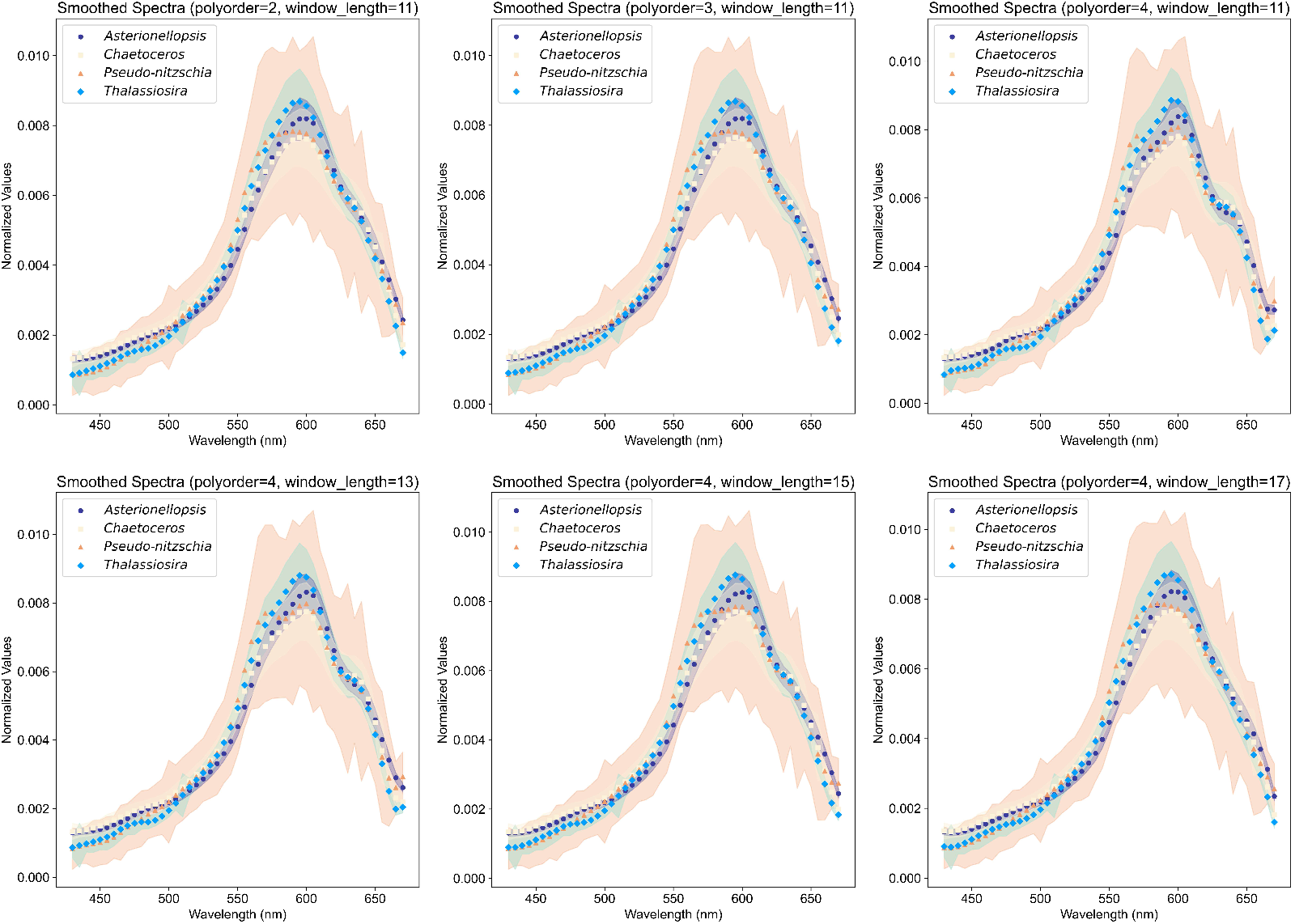
Iterations of the Savitzky-Golay smoothing algorithm applied to the reflectance spectra. The top row shows iterations of the smoothing function varying the polynomial order with a fixed window length of 11 (55 nm). The bottom row shows iterations changing the window-length from 13 (65 nm) to 17 (85 nm) with a fixed polynomial of 4. We selected the combination that preserved the most spectral features while removing noise. The two combinations that did this best were of polynomial size 4, with a window size of 11 or 13. We chose window size 13, matching Vandermeulen et al. 2017.

**Figure 12.**
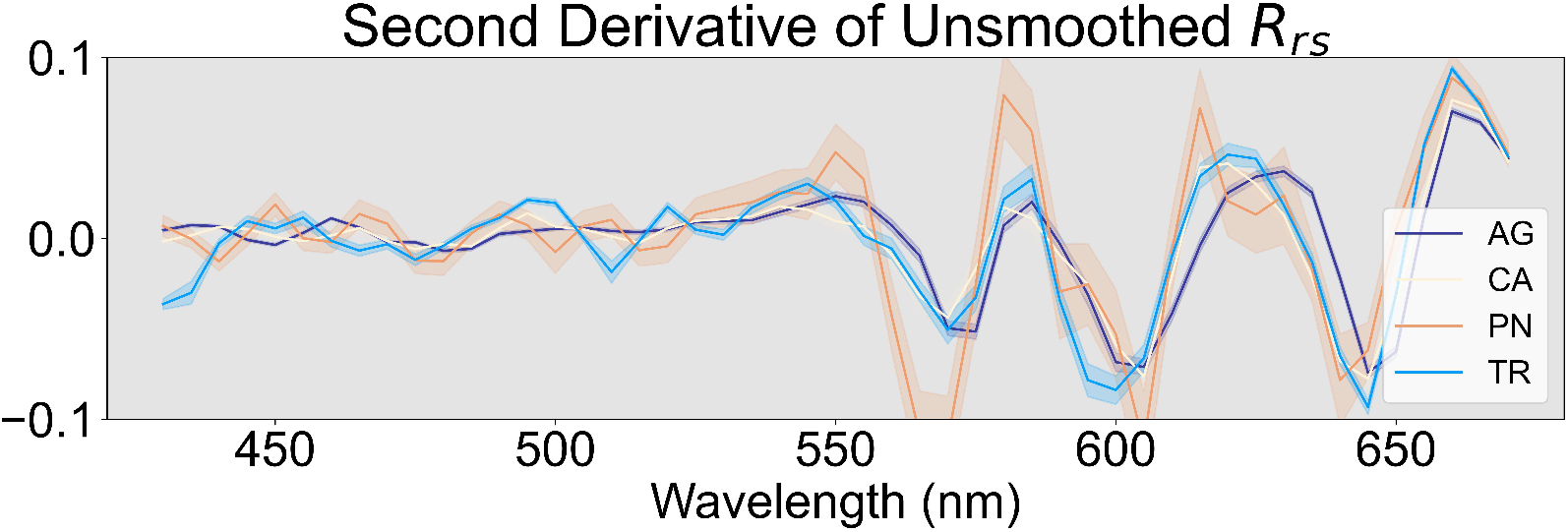
Second derivative of unsmoothed (raw) spectra. One sample taken from each taxon. Note the accentuation of ‘features’ in wavelengths shorter than 550 nm. Whether these are real spectral features that we can glean information from or noise is unknown. Each of these lines are the median values of hundreds to thousands of absorption and backscatter measurements. Noise should be minimal. Yet, smoothing provides a cautionary step to ensure we are not mistaking noise for ‘true’ signal.

**Figure 13.**
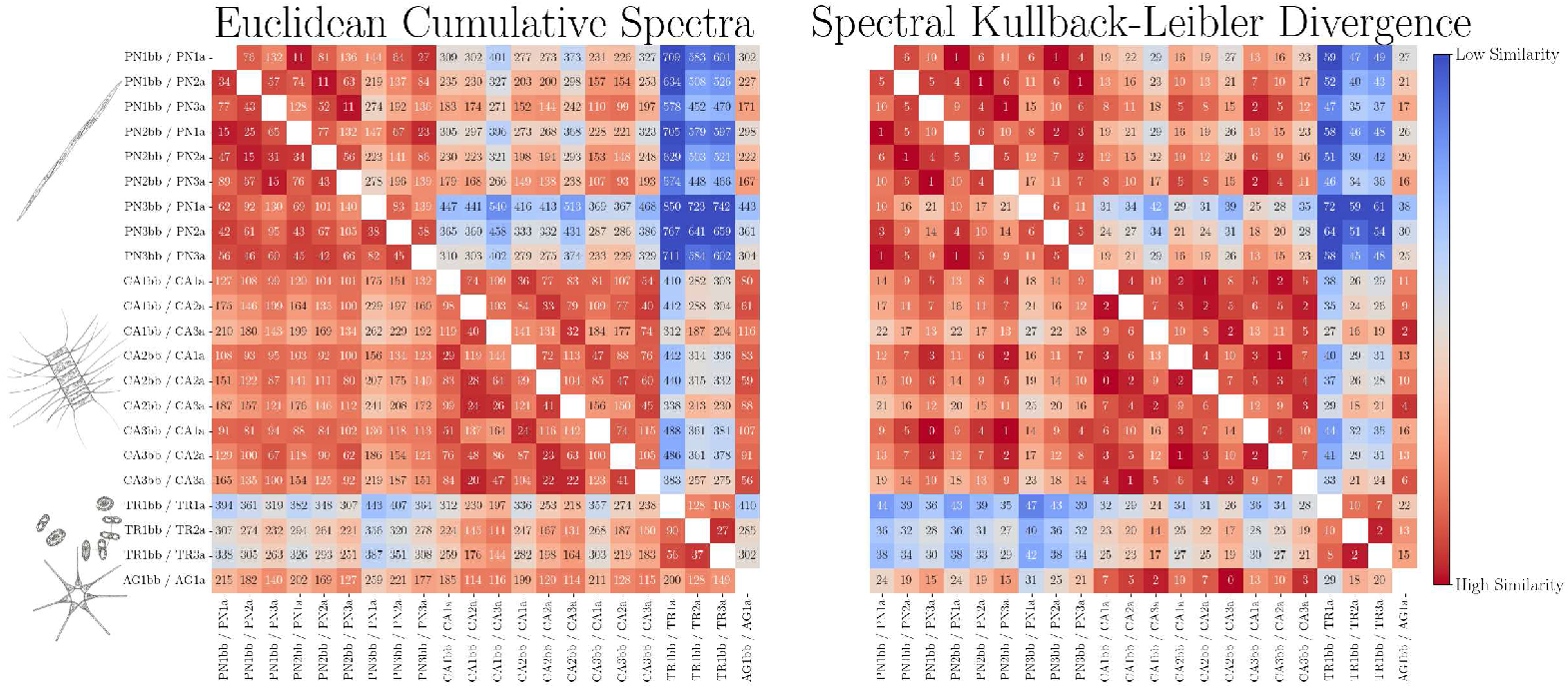
This figure displays two confusion matrices for the ECS and KLPD functions. There is no smoothing or alternate processing. Note the asymmetry in these heatmaps due to differences in integration for ECS (bottom is integrating from blue to red, top is integration from red to blue) and the triangular inequality of KLPD - the triangular inequality means we expect different results based on order of comparison (i.e. spectra a versus spectra b ≠ spectra b versus spectra a). Kullback-Leibler pseudo-divergence performed better when comparing spectra of different diatoms against *Pseudo-nitzschia*, rather than comparing *Pseudo-nitzschia*’s spectra against other diatoms. Euclidean distance of cumulative spectra performs better when integrating the spectra from red to blue wavelengths, rather than blue to red. All values for KLPD and ECS are returned in absolute values since we are interested in the magnitude of the differences and not the sign.

